# Differential ubiquitination as an effective strategy employed by Blood-Brain Barrier for prevention of bacterial transcytosis

**DOI:** 10.1101/2021.06.20.449199

**Authors:** Smita Bhutda, Sourav Ghosh, Akash Raj Sinha, Shweta Santra, Aishwarya Hiray, Anirban Banerjee

**Affiliations:** Bacterial Pathogenesis Lab, Dept. of Biosciences and Bioengineering, Indian Institute of Technology Bombay, Powai, Mumbai 400076, INDIA

**Keywords:** *Streptococcus pneumoniae*, Blood-Brain Barrier, Ubiquitination, K48-Ub, K63-Ub, Autophagy, Proteasomal degradation

## Abstract

The protective mechanisms of blood-brain barrier (BBB) prohibiting entry of blood borne pathogens and toxins into the central nervous system (CNS) is critical for maintenance of homeostasis in the brain. These include various forms of intracellular defence mechanisms which are vital to block bacterial transcytosis, the major route of trafficking adopted by meningeal pathogens to transit into the CNS. However, mechanistic details of the defence mechanisms and their exploitation to prevent bacterial meningitis remain unexplored. In this study, we established that brain endothelium driven ubiquitination acts as a major intracellular defence mechanism for clearance of *S. pneumoniae,* a critical neurotropic pathogen, during its transit through BBB. Our findings suggest that brain endothelium employs differential ubiquitination with either K48 or K63-Ub chain topologies as an effective strategy to target SPN towards diverse killing pathways. While K63-Ub decoration triggers autophagic killing, K48-Ub directs pneumococcus exclusively to the proteasome machinery. Indeed, time lapse fluorescence imaging involving proteasomal marker LMP2 revealed that in BBB, majority of the ubiquitinated SPN were cleared by proteasome. Fittingly, pharmacological inhibition of proteasome and autophagy pathway not only led to exclusive accumulation of K48-Ub and K63-Ub marked SPN, respectively, but also triggered significant increment in intracellular SPN burden. Moreover, genetic impairment of formation of either K48 or K63-Ub chain topology demonstrated that though both chain types play important roles in disposal of intracellular SPN, K48-Ub chains and subsequent proteasomal degradation has more pronounced contribution towards ubiquitinated SPN killing in brain endothelium. Collectively, these observations for the first time illustrated a pivotal role of differential ubiquitination in orchestrating a symphony of intracellular defence mechanisms blocking pathogen trafficking into the brain which could be further exploited to prevent bacterial CNS infections.

**IMPORTANCE:** Among the different cellular barriers present in the human body, Blood-Brain Barrier (BBB) is unique as it not only provides structural integrity but also protects the central nervous system (CNS) from pathogen invasion. In recent past, ubiquitination, which is known to be involved in protein quality control and cellular homeostasis, has been proven to be critically involved in pathogen clearance. In this study, employing *S. pneumoniae* as a model CNS pathogen, we wanted to decipher the critical contribution of ubiquitination in protective mechanism of BBB while tackling bacterial entry into the CNS. Our results suggest, that BBB deploys differential ubiquitination as an effective strategy to prevent neurotropic bacterial trafficking into the brain. By portraying a comprehensive picture of ubiquitin coat on SPN, we figured out that different ubiquitin chain topologies formed on the pneumococcus dictated the selection of downstream degradative pathways, namely, autophagy and proteasomal machinery. Amongst these, contribution of proteasomal system in clearance of pneumococcus was found to be more pronounced. Overall our study revealed how BBB deploys differential ubiquitination as a strategy to trigger autophagy and proteasomal system, which work in tandem to ensure brain’s identity as an immunologically sterile site.

## INTRODUCTION

Human body is composed of multiple cellular barriers that primarily safeguards deeper tissues that are susceptible to infection by pathogens. Among these host cellular barriers, Blood Brain Barrier (BBB) stands out, as this a specialized structural and functional barrier aids maintenance of central nervous system (CNS) homeostasis by blocking entry of toxic metabolites and pathogens circulating in the bloodstream into the CNS, keeping it immunologically sterile (1, 2). Despite the protective architecture of BBB, some neurotropic pathogens such as *Streptococcus pneumoniae* (3), *Neisseria meningitidis* (4), *Haemophilus influenzae* (5), *Escherichia coli* K1 (6) etc. invade brain endothelial cells and breach the BBB transcellularly to cause meningitis. Transcellular invasion into the endothelial cells also provide a safe refuge to the pathogens protecting them from the systemic immunity and facilitating relapse of bacterial infection (7). Due to all these, regardless of advances in antibiotic therapy and vaccines, bacterial meningitis still remains a burning global health issue with high rate of morbidity and mortality (8).

*Streptococcus pneumoniae* (SPN, pneumococcus), the Gram positive α-hemolytic bacteria, is the most common cause of the bacterial meningitis worldwide (9). Employing a plethora of surface proteins, SPN adheres to the brain endothelium and subsequently invades it via various endocytic pathways (10). Following invasion, SPN thwarts intracellular killing by classical endo-lysosomal pathway by secreting a cholesterol dependent pore forming toxin, pneumolysin (Ply). Escape of the SPN from lysosomal killing by virtue of Ply elicits autophagy, facilitating intracellular degradation of SPN and arming the host with an alternate strategy to curb bacterial dissemination. However, in a substantial fraction of intracellular SPN, Ply can pierce the autolysosomal membrane leading to its cytosolic escape (11). Clearance of this cytosolic SPN pool that dodged host intracellular killing pathways is critical to avert bacterial dissemination into the CNS.

Recently, we and others have reported that in brain endothelial cells, cytosol exposed SPN gets ubiquitinated which led to its clearance and limiting its entry into the brain (11, 12). Apart from facilitating pathogen clearance, the ubiquitin system plays a major role in wide range of cellular functions like signal transduction, endosome trafficking, protein sorting, protein degradation etc. (13). The ubiquitin molecule which is attached to the substrate by the action of E1, E2 and E3 conjugating enzymes, can form unique structures on substrates by polyubiquitination, utilizing either the N-terminal methionine residue (M1) or any lysine residue among the eight residues it possess (K6, K11, K27, K29, K33, K48 and K63) (14). The unique ubiquitin topologies thus formed are selectively recognized by various polyubiquitin binding effector proteins (15), subsequently governing downstream signalling events (13).

Although, there are ample evidence suggesting crucial role of ubiquitination in pathogen clearance (11, 16), the mechanistic details regarding how ubiquitination triggers bacterial killing and role of specific chain topologies formed on bacteria and their subsequent contribution towards choice of bacterial killing pathways remain elusive. This is particularly true for brain endothelium where ubiquitination could play a major role in its sentinel function by blocking pathogen entry into the CNS. In this study, using *Streptococcus pneumoniae* as a model pathogen, we for the first time, portray a comprehensive picture of ubiquitination of a pathogen with varying ubiquitin chain topologies during transit through BBB. Our study demonstrated that the differential ubiquitination governing diverse killing pathways, particularly K48-ubiquitination directed proteasomal pathway, is an efficient strategy deployed by the brain endothelium to curb pathogen transit to CNS.

## RESULTS

### Ubiquitination triggers both lysosome dependent and independent clearance of SPN in brain endothelium

Following invasion in brain endothelium and during transcytosis, catastrophic damage to the pneumococcus containing vacuolar (PCV) membrane by the pore forming toxin pneumolysin triggers ubiquitination of SPN. However, mechanisms adopted by the brain endothelial cells to clear such ubiquitinated SPN remain elusive. To determine this, we first tested whether ubiquitinated PCVs associate with lysosomes for subsequent degradation. In classical bacterial killing pathways, bacteria containing endosomes gradually get acidified and finally fuse with lysosomes for ultimate destruction (17). We used Lysotracker as an acidotropic probe to evaluate acidification of the vacuoles (Fig. 1A-B). Surprisingly, confocal microscopy analysis revealed that in brain endothelial cells only a minor subset of the ubiquitinated SPN associated with lysotracker (Ub^+^ Lyso^+^) (35.61 ± 4.9%) while majority (64.38 ± 4.89%) did not get sequestered in acidified vacuoles (Ub^+^ Lyso^-^) (Fig. 1C). Since non-acidification or neutralization of PCVs can occur even after maturation and fusion with lysosomes, due to activity of different bacterial virulence factors including cholesterol dependent cytolysins (18), we evaluated accumulation of Cathepsin B, a lysosomal cysteine protease in the ubiquitinated PCVs (Fig 1D-E). Our results suggest that a significant subset (61.3 ± 2%) of the ubiquitinated PCVs did not accumulate Cathepsin B (Fig. 1F). Simultaneous presence of SPN in acidified lysosome like vacuoles as well as non-acidified compartments imply that brain endothelial cells employ two parallel pathways for clearance of ubiquitinated SPN. While very few SPN is driven towards the lysosome dependent killing, the major subset of ubiquitinated SPN clearance involves a hitherto speculative non-lysosomal pathway.

**Figure 1.**
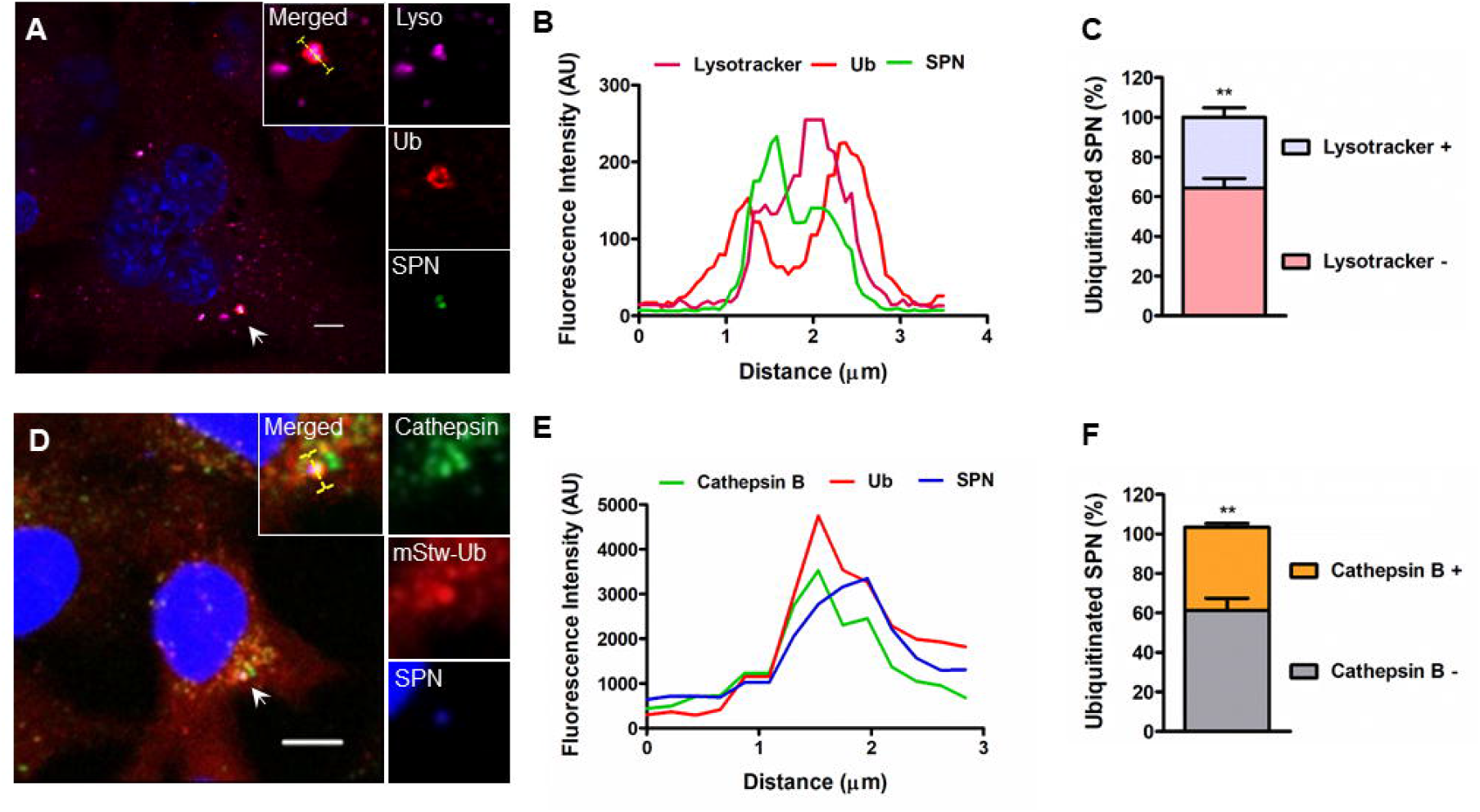
Ubiquitination triggers both lysosome dependent and independent clearance of SPN in brain endothelium. **A.** Confocal micrograph showing association of ubiquitin (red) positive SPN (green) with LysoTracker (pink) at 9 h p.i. hBMECs were infected with GFP expressing SPN and stained with Anti-ubiquitin Ab and acidotropic dye LysoTracker Deep Red. Arrowhead designates bacteria shown in insets. Scale bar, 5 μm. **B.** Fluorescent line scan across the yellow line in the merged inset in “A”. **C.** Quantification of ubiquitinated SPN’s association with LysoTracker in hBMECs at 9 h p.i. n ≥ 100 bacteria per coverslip. Data are presented as mean ± SD of triplicate experiments. Statistical analysis was performed using Students t-test. ** *p* < 0.005. **D.** Confocal micrograph showing association of ubiquitin (red) positive SPN (blue) with lysosomal enzyme cathepsin B (green) at 9 h p.i. hBMECs stably expressing mStrawberry-Ub (red) were infected with SPN and stained with anti-cathepsin B Ab. Arrowhead designates bacteria shown in insets. Scale bar, 5 μm. **E.** Fluorescent line scan across the yellow line in the merged inset in “D”, depicting co-localization of SPN with cathepsin B. **F.** Quantification of co-localization of ubiquitinated SPN with cathepsin B in hBMECs at 3 h p.i. Data are presented as mean ± SD of triplicate experiments. n ≥ 100 bacteria per coverslip. Statistical analysis was performed using Students t-test. ** *p* < 0.005.

### Ubiquitinated SPN is cleared by both autophagy and proteasomal pathway

Given that clearance of majority of ubiquitinated SPN required a lysosome independent cellular pathway, we sought to test the host proteasomal system as an alternate degradative pathway employed by BBB for tackling the invaded SPN. To accomplish this, we generated a transfected hBMEC line stably expressing mStrawberry-Ub and GFP-LMP2, one of the subunits of 26S immunoproteasome by transfecting cells with pMRX-IRES-Puro-GFP-LMP2 and pMRX-IRES-Blast-mStw-Ub plasmids. Time lapse live cell imaging of ubiquitinated SPN labelled with DRAQ5 for elongated period, surprisingly revealed accumulation of GFP-LMP2 to approximately 62.5 ± 2.5% of the ubiquitinated SPN (Fig 2E). We also observed gradual degradation of the ubiquitinated SPN while being associated with GFP-LMP2 (Fig. 2A-B and Movie S1), establishing key role of proteasome in SPN clearance in brain endothelium. Live cell imaging was also performed on a hBMEC cell line stably expressing both mStrawberry-Ub as well as LC3-GFP to decipher contribution of autophagy-lysosome pathway in SPN clearance. We observed that approximately 32.5 ± 3.2% mStrawberry-Ub marked, DRAQ5 labelled SPN, displayed sequestration by LC3-GFP ring like structures with time and eventual degradation while remaining confined within these LC3-GFP structures (Fig. 2C-D and S2 Movie). To further evaluate which process among autophagy and proteasomal pathway is more efficient in tackling intracellular SPN burden during its trafficking through BBB, we treated HBMECs with either 3-MA (autophagy inhibitor) or MG132 (26S proteasome inhibitor) and performed intracellular survival assay. Our results suggest that at non-cytotoxic concentrations (Fig. S1A), both 3-MA or MG132 significantly improved survival ability of SPN in brain endothelium, without affecting internalization, corroborating our time lapse microscopic data (Fig. S1B-D). Intriguingly, inhibiting proteasomal machinery led to 3.8 and 2.5 fold survival advantage to the intracellular SPN in comparison to 2.6 and 1.9 fold survival benefit provided by blocking cellular autophagy at 6 and 12 h p.i., respectively (Fig 2F). Collectively, these results suggest that though both autophagy and proteasomal degradation, are essential for promoting ubiquitinated SPN clearance in brain endothelium, contribution of the proteasomal pathway is more pronounced.

**Figure 2.**
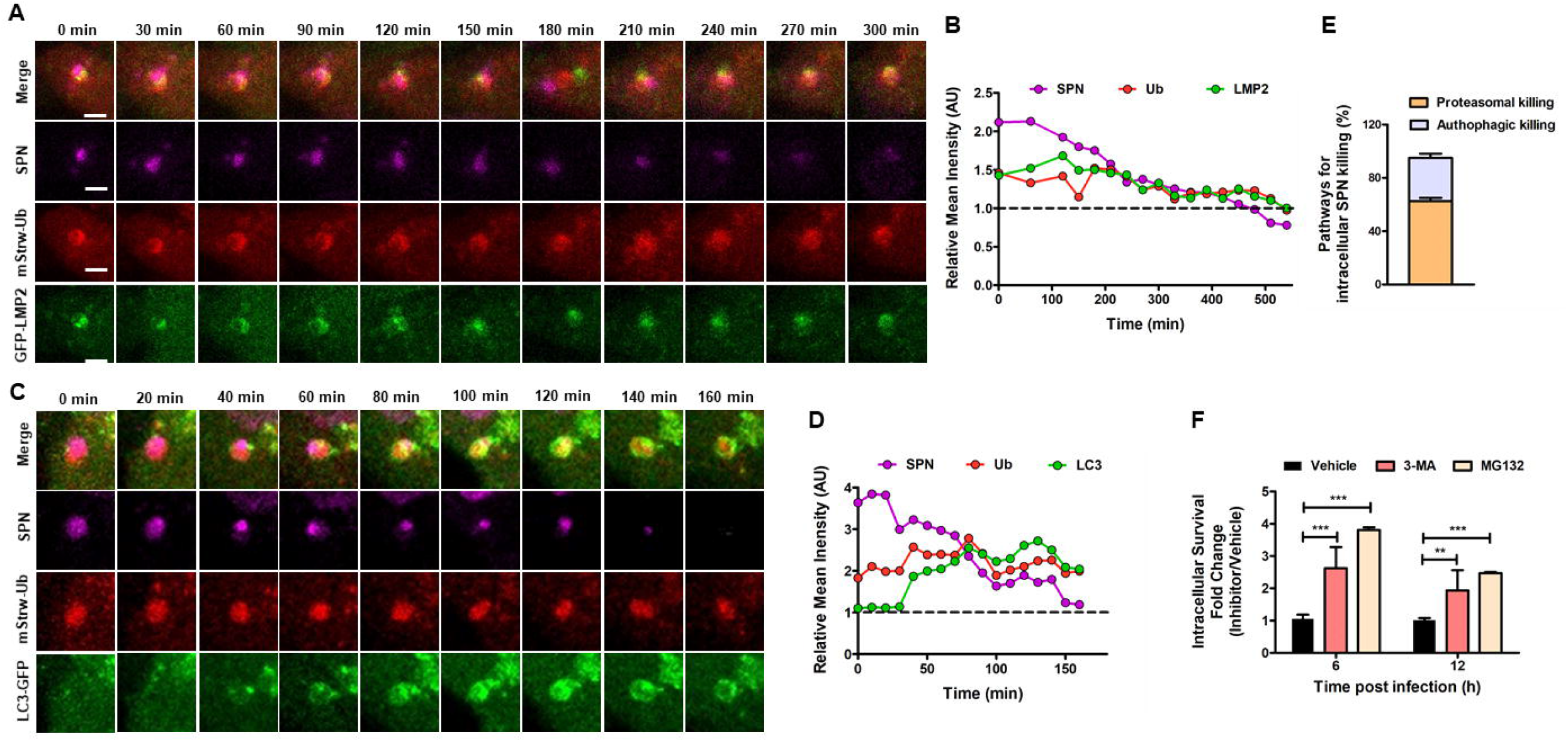
Ubiquitinated pneumococcus is cleared by both autophagy and proteasomal pathway. **A, C.** Representative time-lapse montage for degradation of ubiquitinated SPN in proteasome (A) and autophagy (C) dependent manner. hBMECs stably expressing mStrawberry-Ub (red) and GFP-LMP2 (green) or LC3-GFP (green) were infected with DRAQ5 stained WT SPN (magenta). Live imaging of infected cells was performed for extended hours beginning at 6 h p.i. under a confocal microscope and time-lapse series with intervals of 30 min are represented. The stills in “A” and “C” correspond to movies shown as Movies S1 and S2, respectively. Scale bar, 2 μm. **B, D.** Temporal quantification of ubiquitin, SPN and LMP2 (B) or LC3 (D) fluorescence intensities either relative to fluorescence in the cytosol (Cyt) (for Ubq and LMP2 or LC3) or fluorescence signal of hBMEC nuclei (Nuc) (for SPN). Relative mean intensity for LMP2 or LC3 (I_PCV_/I_Cyt_) in case of ubiquitinated SPN displayed values > 1.0, depicting proteasome (B) or autophagy (D) dependent killing of SPN, respectively. In both cases relative mean intensity values for bacteria (I_SPN_/I_Nuc_) remains > 1.0 for substantially long period of time before finally disappearing. **E.** Percentage of ubiquitinated SPN undergone LC3-dependent or LMP2 mediated killing as examined by time-lapse imaging of hBMECs stably expressing Ubq-mStrawberry and LC3-GFP or GFP-LMP2, respectively. n = 50. **F.** Comparison of intracellular survival of SPN in hBMECs following pre-treatment with either 3-MA (1 mM) or MG132 (10 μM) or DMSO (vehicle). Fold change in intracellular survival at indicated time points post infection is calculated by multiplicity of percent SPN survived (relative to 0 h) in presence of 3-MA or MG132 in comparison to DMSO. Data are presented as mean ± SD of triplicate experiments. Statistical analysis was performed using two-way ANOVA (Bonferroni test). ** *p* < 0.005, \*\*\**p* < 0.001.

### Differential ubiquitination of SPN with varied chain topologies governs choice of host degradative pathways

Since specific ubiquitin chain topologies formed on the substrate dictate downstream signalling pathways triggering precise degradation pathways, we aimed to decipher the composition of ubiquitin coat covering intracellular SPN. We hypothesized that differential ubiquitin coat could be the decisive factor in shunting the ubiquitinated SPN towards autophagy or proteasomal pathway in brain endothelium. To accomplish this intracellular SPN was stained with different ubiquitin linkage specific antibodies. Among the different ubiquitin linkage topologies, K48-Ub and K63-Ub chains are primary signal for degradation (19, 20). Interestingly, we observed that brain endothelium decorates SPN with both types of ubiquitin chain topologies (Fig. 3A-D). However, quantification of such events at various time points post infection revealed significant difference in the kinetics of their contribution towards total ubiquitinated SPN, especially at later stages of infection. At 6 h p.i., K63-Ub SPN had reduced drastically from 68.15 ± 6.25% to 43.38 ± 4.81%, and subsequently to 31.54 ± 3.33% at 9 h and finally to 17.9 ± 3.69% at 12 h p.i. However, unlike K63-Ub chains, contribution of K48-Ub chains to the total ubiquitinated SPN remained fairly constant till extended time of infection (70.87 ± 1.94%, 67.87 ± 5.67%, 64 ± 2% and 73.64 ± 3.18% at 3, 6, 9 and 12 h p.i., respectively) (Fig. 3E). This could be due to the fact that with increased time, progressively larger damage to the PCV membrane could lead to higher degree of decoration with K48-Ub chain topology. Alternatively, faster degradation kinetics of K63-Ub chain types could have triggered its drastic reduction with time.

**Figure 3.**
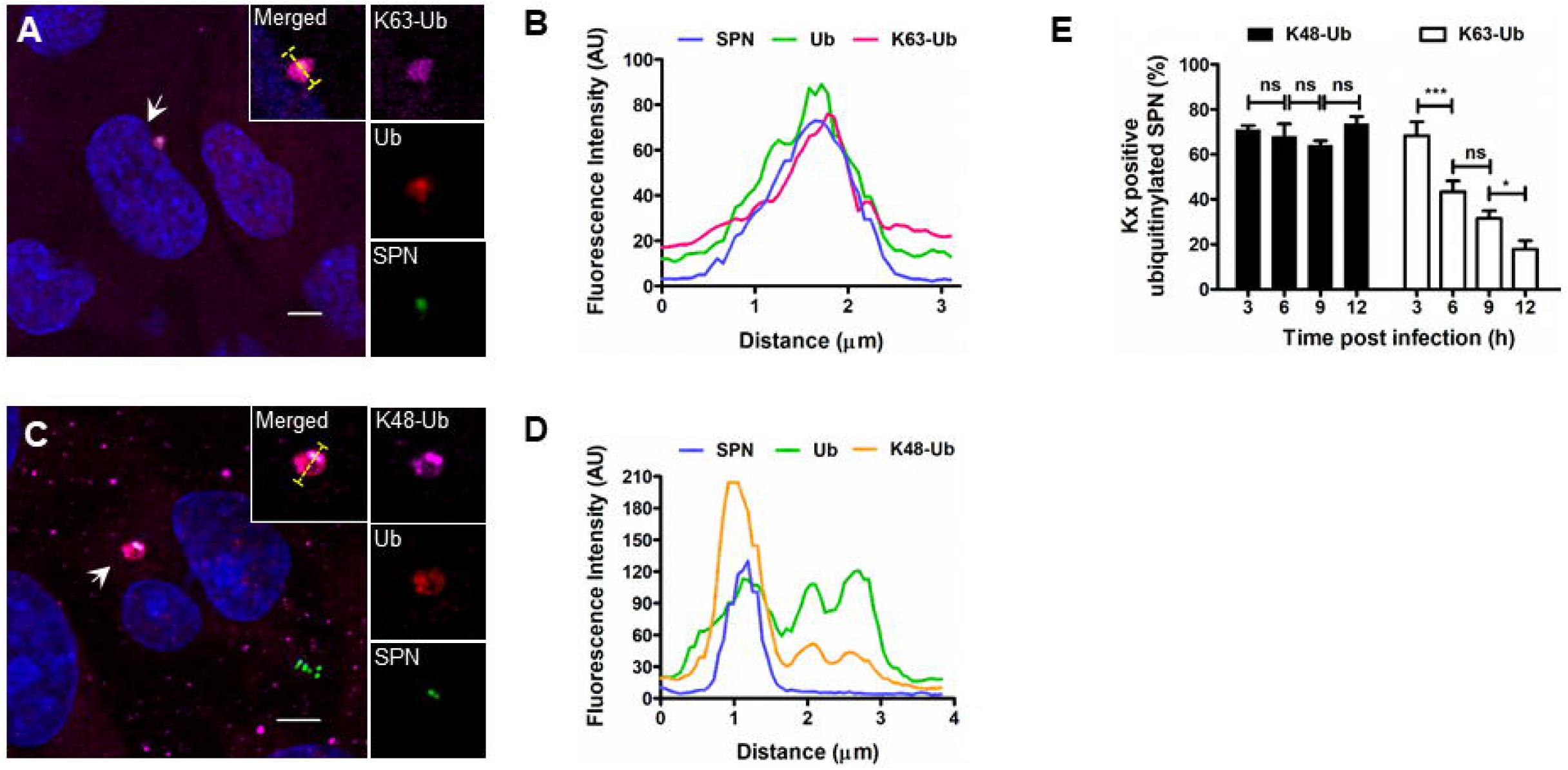
SPN is ubiquitinated with different ubiquitin chain topologies in brain endothelium. **A, C.** Confocal micrographs showing association of ubiquitinated (red) SPN (magenta) with specific ubiquitin chain types (K48 in “A” and K63 in “B”) at 6 h p.i. hBMECs stably expressing mStrawberry-Ub was infected with SPN and stained with Anti-K48-Ub (A) or Anti-K63-Ub (C) linkage specific Abs. Arrowhead designates event shown in insets. Scale bar, 5 μm. **B, D.** Fluorescent line scan across the yellow line in the merged inset in “A and C” are shown in “B” and “D”, respectively. **E.** Contribution of K48-Ub and K63-Ub decorated SPN towards total ubiquitinated SPN at 3, 6, 9 and 12 h p.i. n ≥ 100 ubiquitinated bacteria per coverslip. Data is represented as mean ± SD of triplicate coverslips. Statistical analysis was performed using one-way ANOVA (Tukey’s multiple comparison test). ns, non-significant, \**p* < 0.05, \*\*\**p* < 0.001.

We then aimed to determine the role of K63-Ub and K48-Ub chains in driving the intracellular SPN towards autophagy or proteasomal system, the two established pathways involved in clearance of ubiquitinated intracellular pathogens. LC3-GFP and GFP-LMP2 were used as markers for autophagy and proteasomal system, respectively. We observed clear association of LC3 with K63-Ub marked SPN (Fig. 4C-D and S2C-D) while K48-ubiquitinated SPN exhibited exclusive accumulation of GFP-LMP2 signal (Fig. 4A-B and S2A-B). Quantification of such events revealed that majority of K48-Ub marked SPN (57.33 ± 2.51%) was found to be associated with GFP-LMP2, while a minor population (9.32 ± 2.3and 0%) exhibited co-localization with LC3-GFP (Fig. 4E). Further, MG132 treatment to the hBMECs to impair proteasomal activity caused accumulation of the K48-Ub SPN without affecting degradation of K63-Ub positive SPN (Fig. 4G), implying that in brain endothelium K48-Ub drives SPN towards proteasomal killing. On the contrary, 45.76 ± 3.28% of the K63-Ub SPN population was found to be co-localized with LC3-GFP, while only 8.33 ± 1.52% showed association with GFP-LMP2 (Fig. 4F). Additionally, treatment of hBMEC with 3-Methyladenine led to 2 fold significant increase in the K63-Ub marked SPN and showed no effect on the K48-Ubiquitinated SPN (Fig. 4G). These observations collectively demonstrate that brain endothelial cells employ K63-Ubiquitination to preferentially target intracellular SPN towards autophagic killing by sequestering it inside LC3 marked autophagosome, while K48-Ubiquitination promotes SPN killing via proteasomal machinery.

**Figure 4.**
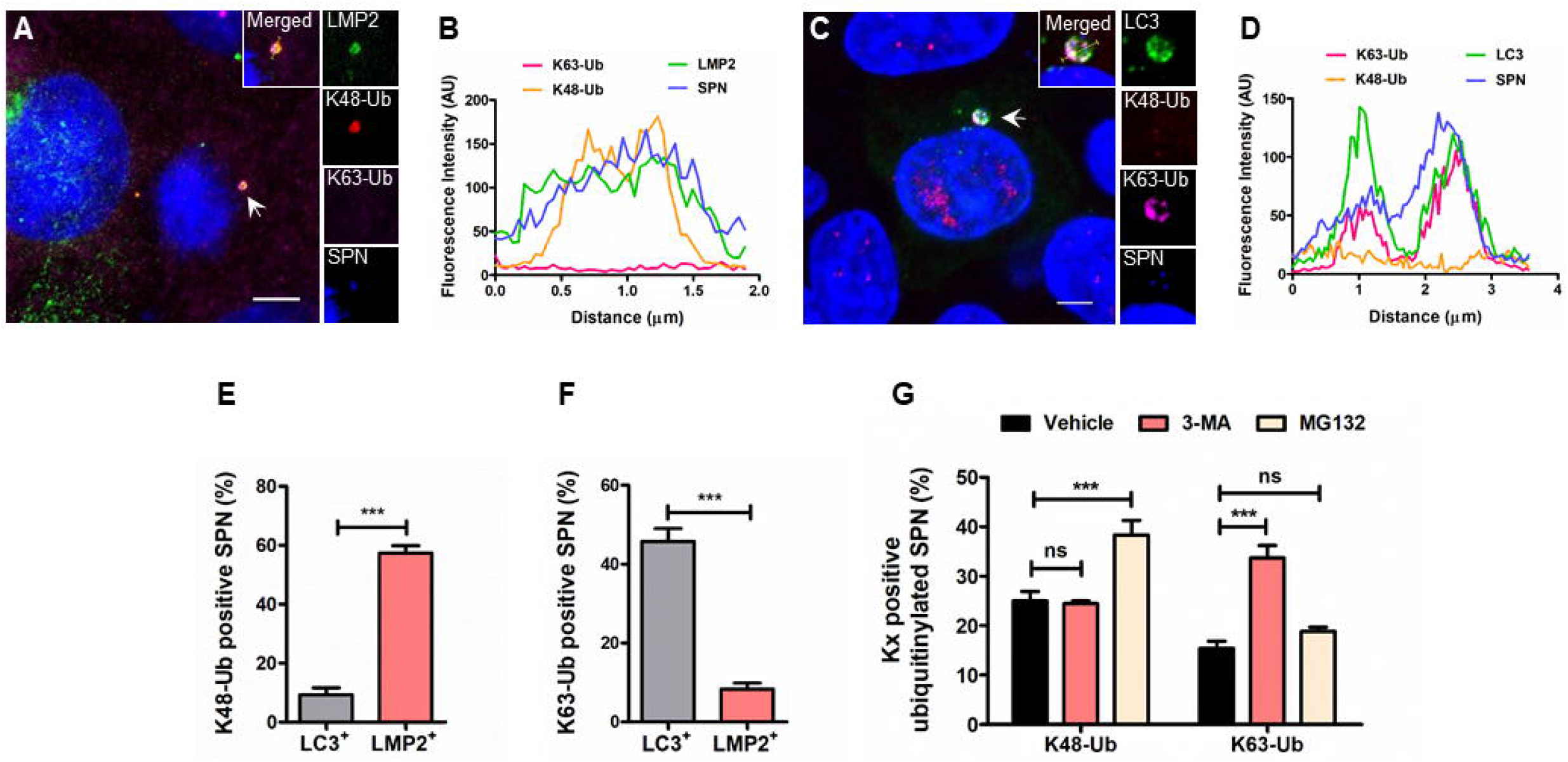
Differential ubiquitination of SPN with varied chain topologies governs choice of host degradative pathways. **A, C.** Confocal micrographs showing association of K48-Ubiquitinated (red) SPN (blue) with LMP2 (green) (A) or K63-Ubiquitinated (magenta) SPN (blue) with LC3 (green) (C) in hBMEC at 9 h p.i. hBMECs stably expressing GFP-LMP2 (A) or LC3-GFP (C) were infected with Hoechst stained SPN and stained with Anti-K63-Ub and Anti K48-Ub linkage specific Abs. Arrowhead designates event shown in insets. Scale bar, 5 μm. **B, D.** Fluorescent line scan across the yellow line in the merged inset in “A” and “C” is shown in “B” and “D”, respectively. **E, F.** Quantitation of association of LC3 and LMP2 with K48-Ub (E) or K63-Ub (F) positive SPN. n ≥ 50 K48/K63-ubiquitinated bacteria per coverslip. Data is represented as mean ± SD of triplicate hBMEC cultures. Statistical analysis was performed using unpaired Student’s t test. *** *p* < 0.001. **G.** Quantification of SPN ubiquitinated with K48-Ub or K63-Ub chain topologies, at 9 h p.i. following pre-treatment of hBMECs with 3-MA (1 mM) or MG132 (10 μM). n ≥ 50 K48/K63-ubiquitinated bacteria per coverslip. Data is represented as mean ± SD of triplicate coverslips. *p*-values are from one-way ANOVA analysis with Dunnett’s multiple comparisons test. ns, non-significant; *** *p* < 0.001.

### K48-Ub chains contributes primarily for controlling intracellular SPN burden in brain endothelium

After deciphering the composition of the ubiquitin coat of PCVs and their individual role, we next investigated the comparative contribution of different ubiquitin chain topologies in promoting clearance of intracellular SPN during its passage through the brain endothelium. To accomplish this, we inhibited formation of K48 and K63 ubiquitin chains individually in hBMECs by expressing Ub^K48R^ and Ub^K63R^ mutant ubiquitins from pLKO-Puro-tet-Ub^K48R^-GFP and pLKO-Puro-tet-mStrawberry-Ub^K63R^ plasmids, respectively. Given that ubiquitination is indispensable for maintenance of cellular homeostasis and both stable expression of mutant ubiquitin or deletion of WT ubiquitin is lethal for host cell, we expressed mutant ubiquitins under tetracycline inducible promoter using a Tet-ON/OFF system in brain endothelial cells. This ensured that only in presence of tetracycline, cells expressed mutant Ub (Ub^KxR^) along with Ub^WT^ which is expressed from the genome. We hypothesized that simultaneous expression of Ub^WT^ and Ub^KxR^ in hBMECs would prematurely thwart polyubiquitin chain formation due to incorporation of Ub^KxR^ in growing polyubiquitin chain by dominant negative effect. The aborted ubiquitin chain topology therefore formed can not dictate downstream signaling, leading to inefficient degradation of the cargo and its subsequent accumulation (Fig. 5A). Visualization of hBMECs transfected with fluorescently labelled mutant ubiquitins following treatment of non-cytotoxic concentration of tetracycline (Fig. S3A) under confocal microscope showed similar expression of Ub^K48R^-GFP or mStrawberry-Ub^K63R^ or mStrawberry as a control (Fig. 5B). Expectedly, hBMECs expressing Ub^K48R^-GFP and mStrawberry-Ub^K63R^ not only blocked formation of K48-Ub and K63-Ub chain topologies, respectively, but did not perturb the formation of opposite ubiquitin chain types (Fig. 5C). Critically, hBMECs expressing mutant ubiquitin variants (Ub^K48R^-GFP and mStrawberry-Ub^K63R^), though did not show any defect in pneumococcal invasion (Fig. S3B), they were unable to restrict intracellular SPN as efficiently as hBMECs with only Ub^WT^ (Fig. 5D), confirming vital role of ubiquitination in clearance of intracellular SPN in brain endothelium. Further, our findings suggest that though impairment of K63-Ub chain formation promoted SPN survival by a modest margin (1.85 fold), prevention of decoration with K48-Ub chains significantly improved intracellular survival ability of the SPN by 3.91 fold compared to Ub^WT^ expressing hBMEC (Fig. 5D). These findings collectively corroborate with our other observations and unambiguously demonstrate that K48-Ubiquitination and subsequent targeting towards proteasomal killing is an effective strategy adopted by brain endothelium to control intracellular SPN pool.

**Figure 5.**
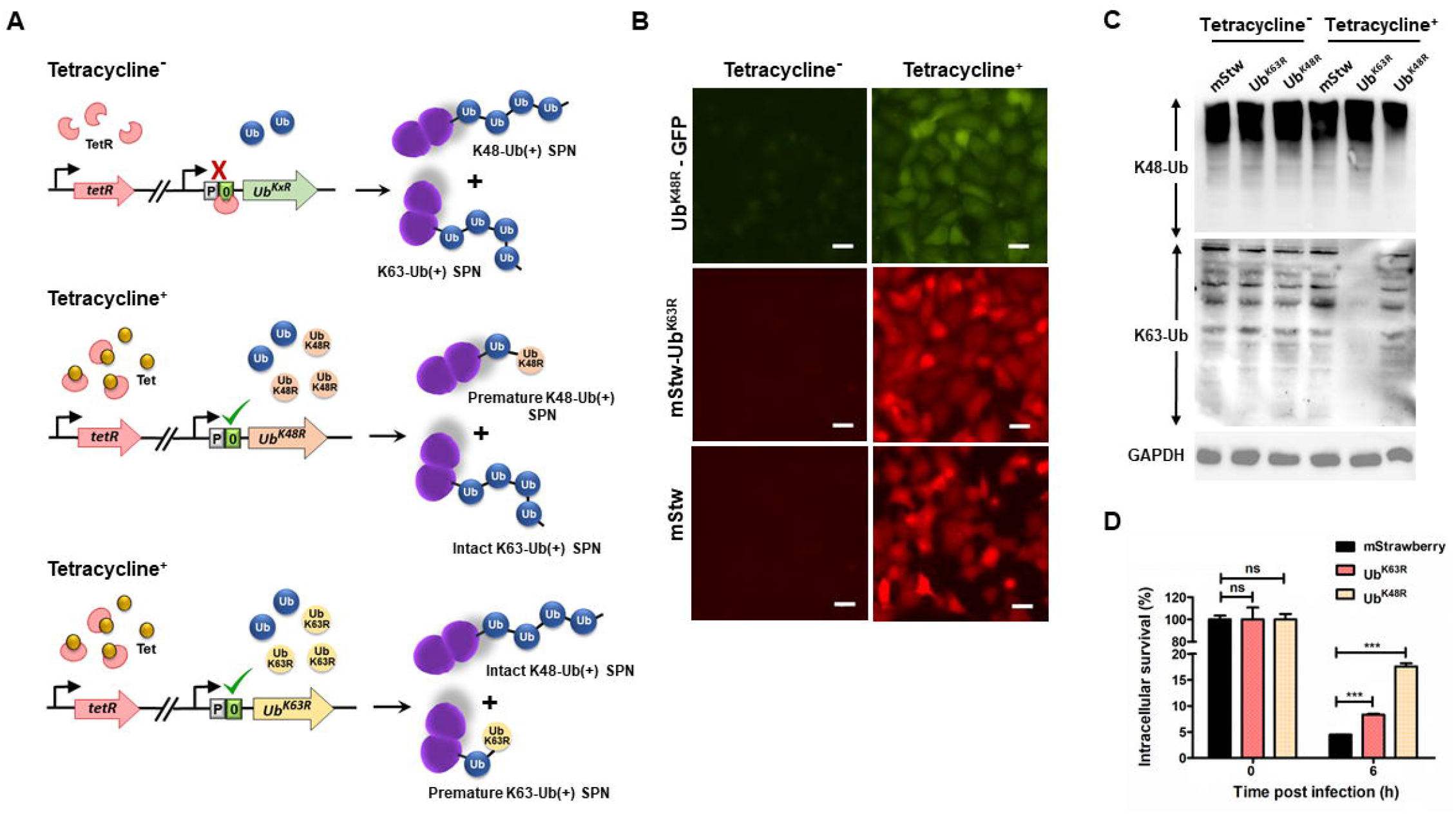
K48-Ub chains contribute primarily for controlling intracellular SPN burden. **A.** Schematic representation of inhibition of SPN decoration with K48-Ub or K63-Ub chain topologies by expressing Ub^K48R^ and Ub^K63R^ mutant ubiquitins under tetracycline inducible promoter. Incorporation of Ub^K48R^ and Ub^K63R^ variants in a growing polyubiquitin chain would thwart further elongation and subsequent signalling, affecting subsequent degradation of substrates by chain specific pathways. **B.** Confocal micrographs depicting hBMECs expressing Ub^K48R^-GFP or mStrawberry-Ub^K63R^ or mStrawberry following transfection with pLKO-Puro-tet-Ub^K48R^-GFP, pLKO-Puro-tet-mStrawberry-Ub^K63R^ and pLKO-Puro-tet-mStrawberry plasmids, respectively, in presence or absence of tetracycline (40 μg/ml). Scale bar, 5 μm. **C.** Evaluation of inhibition of K48-Ub or K63-Ub chain topology formation in hBMECs transfected with either Ub^K48R^-GFP or mStrawberry-Ub^K63R^ mutant variants in presence of tetracycline following immunoblotting with Anti-K48-Ub or Anti-K63-Ub linkage specific Abs. GAPDH served as a loading control. **D.** Intracellular survival efficiency of SPN in hBMECs expressing Ub^K48R^-GFP or mStrawberry-Ub^K63R^ mutant variants at 6 h were calculated as percentage survival relative to 0 h. Data are presented as mean ± SD of triplicate experiments relative to 0 h. Statistical analysis was performed using two-way ANOVA (Bonferroni test). ns, non-significant, *** *p* < 0.001.

## DISCUSSION

Bacterial meningitis occurs because of uncontrolled bacterial replication in the CNS following invasion of the blood brain barrier (BBB) (21). However, the brain endothelium functions as a sentinel to thwart easy passage of pathogens across BBB, protecting the CNS from bacterial invasion. One of the major mechanism employed by the BBB to block bacterial trafficking is endo-lysosomal killing. Though certain neurotropic pathogens have devised strategies to efficiently evade the endo-lysosomal pathway, the brain endothelium adopts alternate means present in its arsenal to clear these escaped pathogens, inhibiting BBB breach. *Streptococcus pneumoniae,* the most prominent meningitis causative microorganism, has been shown to rupture endosomal membrane during transit through BBB due to expression of the pore forming toxin, pneumolysin. This blocks endosome maturation and subsequent lysosomal fusion. Our previous findings demonstrate that brain endothelium ubiquitinates a large subset of such intracellularly persistent SPN for eventual killing (11). In this study, we extended our findings to reveal the details of different molecular mechanisms adopted by brain endothelial cells to dispose such ubiquitinated SPN by decorating them with varying ubiquitin chain topologies (Fig. 6). This lowers pneumococcal burden in BBB following bacterial invasion and prevents their access to central nervous system for establishment of infection.

**Figure 6.**
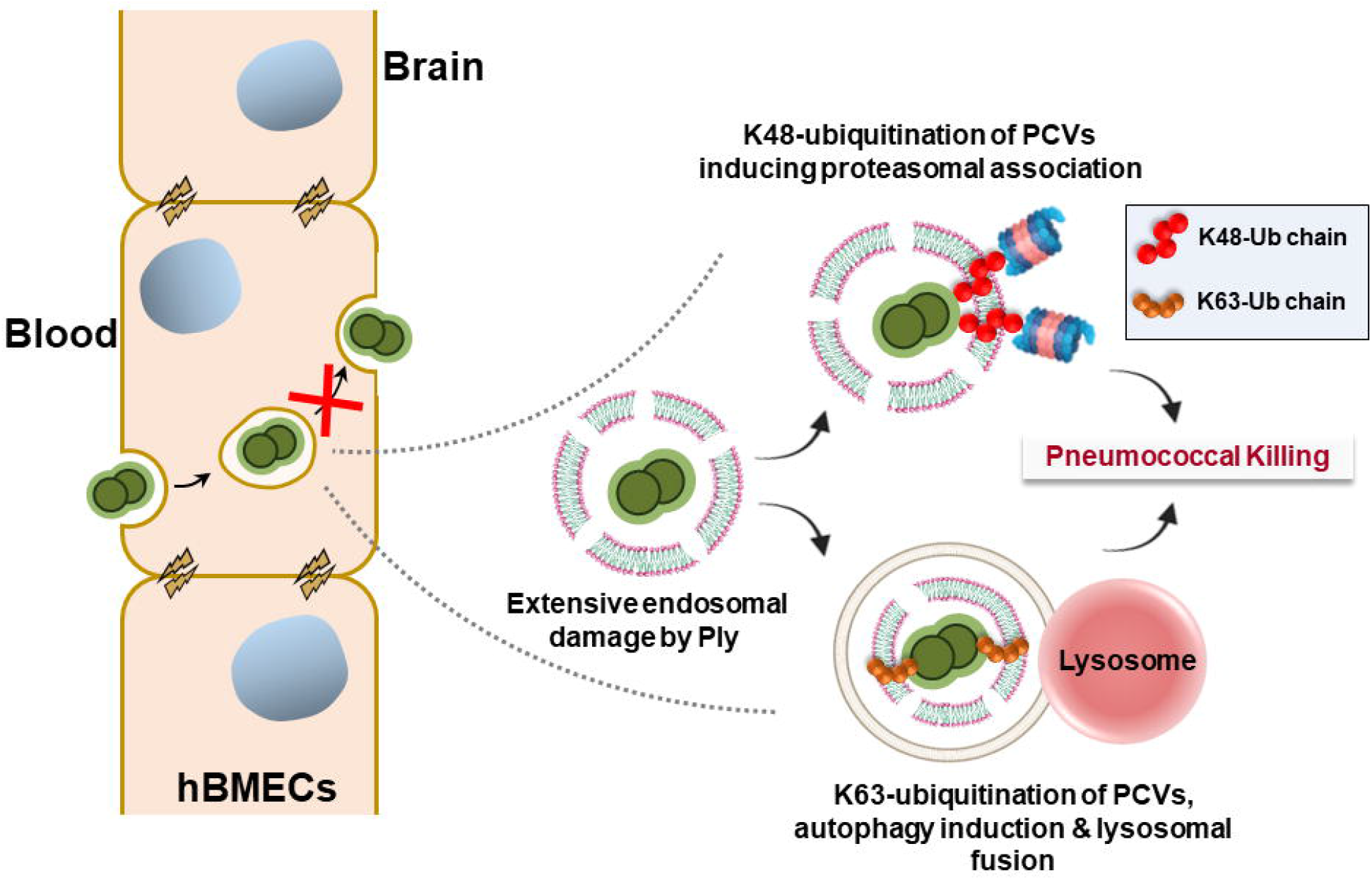
Model depicting differential ubiquitination dependent intracellular clearance of SPN during transit through Blood-brain barrier. Following internalization in brain endothelium, secretion of Ply triggers extensive damage to PCV membrane. This lead to ubiquitination of the PCVs with varying chain topologies. K63-ubiquitination results in sequestration and eventual degradation of SPN in autophagosomes following fusion with lysosomes. On other hand, K48-ubiquitination selectively triggers association and subsequent killing while being associated with proteasomes. A coordinated action of these distinct intracellular defense mechanisms ensures fruitful blockage of SPN transcytosis across brain endothelium, offering protection to the CNS.

In accordance with intracellular fate of other pathogens (22), we also observed acidification of a subset of ubiquitinated PCVs which ultimately fused with lysosomes for degradation in brain endothelium. This was expected as several host autophagy receptors have ubiquitin binding domain to sequester the cargo, including pathogens, in autophagosomes for eventual fusion and degradation in lysosomes. Activation of anti-pneumococcal autophagy was earlier reported in lung epithelial and brain endothelial cells (11, 23, 24). However, we could demonstrate that the specificity for such process is provided by K63-Ub chain types formed on PCVs. Intriguingly, the major population of ubiquitinated PCVs in BBB were neither acidified nor displayed characteristic lysosomal markers. Intracellular bacteria like *Streptococcus pyogenes, Streptococcus pneumoniae, Listeria monocytogenes* etc. which produces cholesterol dependent cytolysins such as Streptolysin O (SLO), Pneumolysin (Ply), Listeriolysin O, respectively, have been reported to rupture not only endosomal but also autophagosomal membrane by virtue of their pore forming ability (11, 25, 26). Progressively larger pores formed by these toxins triggers escape of ubiquitinated pathogens from autophagosomes. But such ubiquitinated autophagosome-escaped pathogens must be cleared by other mechanisms before they transcytose through BBB. Coexistence of a large fraction of such non-lysosomal, ubiquitinated SPN population propelled us to speculate that apart from autophagy, brain endothelium must deploy a parallel degradative pathway for clearance of ubiquitinated intracellular SPN. Our findings reveal that indeed the host proteasomal pathway contributes significantly in intracellular ubiquitinated SPN clearance. Though proteasomal involvement in bacterial killing is proposed previously for *M. tuberculosis* (27), we for the first time, using time lapse fluorescence imaging showed degradation of ubiquitinated bacteria in close association with proteasomal machinery (Figure 2). Indeed, association of proteasome with SPN is reported in mouse brain tissue (12). This further substantiates our claim for involvement of proteasomes as a major mechanism deployed by brain endothelium to restrict pneumococcal passage across BBB, which could be extrapolated to other neurotropic pathogens.

To unravel the factors governing the specificity of the bacterial degradation pathways, we then explored for the contribution of the ubiquitin chain topologies formed on the PCVs. Ubiquitin chain types formed on the substrate which includes cellular proteins, organelles and even bacteria, are reported to govern the choice of further clearance pathways (28). The selectivity of the ubiquitin binding adaptor proteins associated with either autophagy or proteasome dictate the choice of pathways. High local concentrations of K63-linked poly ubiquitin chains acquiring an extended shape preferentially binds to the autophagic adaptors p62, NBR1 and NDP52 driving the cargo towards autophagy (29). In the contrary, K48-linked ubiquitin polymers adopt a compact conformation, favouring a sandwich like binding by the ubiquitin binding adaptor proteins associated with proteasome (30). Our findings for the first time demonstrated a comprehensive composition of the ubiquitin coat of intracellular *S. pneumoniae* involving K48 an K63-Ub chain topologies (Fig. 3). We envisage that this composition of ubiquitin coat on pneumococcus could be of similar order in any host cell type. Difference in the formation kinetics of K48-Ub and K63-Ub suggests that though at early stages of infection PCV bound SPN gets ubiquitinated with different types of ubiquitin topology, at prolonged infection intervals a large subset of SPN is present in cytosol without the shield of any membrane remnants. This is supported by progressive steep decrease in the association of SPN with K63-Ub chains. Since majority of K63-Ubiquitin signal was earlier observed to be associated with PCVs membrane (24), it could be projected that K63-Ub signal is left behind along with damaged endosome remnant when SPN escaped to host cytosol.

In contrary to a previous report that showed reduction in the abundance of K63-Ub SPN in absence of autophagy (24), our findings involving recruitment of LC3-positive autophagosome to the K63-Ub positive SPN coupled with accumulation of K63-Ub positive SPN only in autophagy inhibitor treated cells unambiguously proves that K63-Ub directs SPN towards autophagic killing. This is in accordance with other intracellular pathogens like *M. tuberculosis* that has also been reported to be targeted towards autophagy following ubiquitination with K63-Ub chains by E3 ligase Parkin (22). Critically, exclusive association of LMP2 with K48-Ub positive SPN and further accumulation of K48-Ub SPN when proteasomal machinery of the brain endothelium was impaired established that K48-Ub is responsible for the proteasomal killing of the SPN. Though previously it was observed that K48-Ub SPN accumulates in Atg5 KO MEFs (24), we, however, did not observe any effect of autophagy inhibition on the abundance of K48-Ub positive SPN in hBMECs. We argue that this association of K48-Ub positive SPN with autophagy could be a specific phenotype in MEFs as in majority of the host cells K48 ubiquitination maintains protein homeostasis by directing them to the proteasome (31)

Numerous pathogens like *M. tuberculosis, Salmonella* etc. have been shown to be decorated with K63, K48 and M1-ubiquitin chain topologies, respectively (22, 24, 27, 32). However, relative contribution of individual chain types in the fate of intracellular pathogen have remained elusive. Along with linkage type, length of the chain plays crucial role dictating further ubiquitin mediated signals (33). Insufficient length of the ubiquitin polymer impairs association of the ubiquitin binding adaptor proteins, arresting further degradation of the cargo (34). We targeted this attribute of ubiquitin chain length to compare the impact of each ubiquitin chain types on SPN degradation. Adopting a novel approach to impair formation of one specific type of ubiquitin chains (either K48 or K63), we demonstrated that though both K48 and K63 ubiquitin chains play important roles in clearance of intracellular SPN, K48-Ub chains and subsequent proteasomal degradation has more telling contribution towards ubiquitinated SPN killing in brain endothelial cells. Interestingly, unlike SPN, increase in intracellular *M. tuberculosis* load is similar in absence of K48 and K63 ubiquitin chains (27) implying that contribution of each ubiquitin chain type towards clearance of bacteria varies for different intracellular pathogen.

Collectively, these observations for the first time illustrated a symphony of ubiquitin mediated intracellular defense mechanisms deployed by the blood brain barrier to prevent pathogen replication. Synergy between such defense mechanisms allow the brain endothelium to function as a sentinel for protection of CNS from bacterial entry and maintenance of homeostasis.

## MATERIALS AND METHODS

### Cell culture and transfections

Human brain microvascular endothelial cells (hBMECs, provided by Prof. KS Kim, Johns Hopkins University, USA) (35) were used as model for the BBB. hBMECs have been extensively used for studying interaction of host with meningeal pathogens and characterize host response (36–40). hBMECs are routinely cultured in RPMI 1640 medium (Gibco) supplemented with 10% Fetal Bovine Serum (Gibco), 10% Nu-Serum (Corning), and 1% Non-Essential Amino acids (Gibco) at 37°C in 5% CO_2_. Stably transfected hBMECs overexpressing LC3-GFP, GFP-LMP2 and mStrawberry-Ub fusion proteins were constructed by transfecting hBMECs with pBABEpuro LC3-GFP vector (Addgene, 22405), pMRX-IRES-GFP-LMP2 (GFP-LMP2 was subcloned from pCND3-GFP-LMP2, a kind gift from Prof. J. Neefies, The Netherlands Cancer Institute, The Netherlands) (42) and pMRX-IRES-mStrawberry-ubiquitin (43) (kind gift from Prof. T Yoshimori, Osaka University, Japan). Ub^K48R^-GFP (44) (kind gift from Prof. D. Gray, Ottawa Regional Cancer Centre, Canada), mStrawberry-Ub^K63R^ (45) (kind gift from L. Penengo, University of Zuric-Vetsuisse, Switzerland) and mStrawberry were stably expressed in hBMECs under Tetracycline inducible promoter in pLKO-Puro-tet plasmid (Addgene, 21915). All transfections were performed using Lipofectamine 3000 reagent (Invitrogen) and selections were done in the presence of 2 μg/ml puromycin (Sigma Aldrich) and 2 μg/ml blasticidin hydrochloride (HiMedia) where necessary. For inhibition studies, hBMECs were pre-treated with 3-Methyladenine (1 mM, 24 h) or MG132 (10 μM, 1 h) before infection with bacteria. The concentrations used for 3-MA and MG132 are non-cytotoxic as found out by MTT assay. For induction to express mutant ubiquitin variants (Ub^K48R^-GFP and mStrawberry-Ub^K63R^), cells were treated with freshly prepared tetracycline (40 μg/ml, 15 h) prior to the assay and formation of ubiquitin variants were verified using immunoblotting.

### Bacterial strains and constructs

SPN strain R6 (serotype 2, unencapsulated) was obtained from Prof. TJ Mitchell (Univ. of Birmingham, UK). SPN was routinely grown in Todd-Hewitt broth (THB) supplemented with 1.5% yeast extract (THY media) at 37□C in 5% CO_2_. To construct Tetracycline resistant SPN, *tet*^R^ gene was amplified from pYPQ132D (Addgene, 134348) and cloned in between XbaI and XhoI and sites of the shuttle vector pIB166 (46) (kind gift from Prof. Indranil Biswas, Kansas University Medical Centre, USA) and transformed into wildtype SPN strain using competence stimulating peptide 1 (100 ng/ml). Tet^R^-SPN was further used to infect mutant ubiquitin expressing cell lines as mentioned earlier.

### Plasmids, antibodies and reagents

Fluorescent SPN was developed by transformation of WT SPN with *hlpA-GFP* fusion cassette (kindly provided by Prof. JW Veening, University of Groningen, Netherlands). The following primary antibodies were used in this study. Cathepsin B (Abcam, ab58802), anti-Ubiquitin K48-Specific (Merk Millipore, 05-1307), anti-Ubiquitin K63-Specific (Enzo Life science, BML-PW0600-0025), Anti-Ubiquitin (Enzo Life Sciences, BML-PW-8810), GAPDH (Millipore, MAB374). Fine chemicals used in this study include 3-Methyladenine (M9281) and MG132 (474791), from Sigma Aldrich; Lysotracker Deep Red (Invitrogen, L12492) and DRAQ5 Deep red (BD Pharmingen, 564903).

### Penicillin gentamicin protection assay

Infection assays were performed as described earlier (47). Briefly, fully confluent hBMEC monolayers in 1% collagen coated 24 well plate was infected with SPN (O.D.600 ~ 0.4) at a multiplicity of infection (MOI) of 10 for 1 h. Following infection, extracellular SPN were killed by incubating infected cells for 2 h in culture medium containing penicillin (10 μg/ml) and gentamicin (400 μg/ml). After several washes, cells were trypsinized (0.025% trypsin EDTA), lysed (0.025% Triton X-100) and the lysates were spread plated on BHI (Brain Heart Infusion) agar plates for enumeration of bacterial colonies. Bacterial invasion efficiency was calculated as (recovered CFU/initial inoculum CFU) × 100%. To assess intracellular survival of bacteria, cell lysates were prepared similarly and spread plated at indicated time intervals following 1 h of infection and 2 h of antibiotic treatment. Surviving bacteria at different time points were enumerated and was represented as percent survival at indicated time points relative to 0 h.

### Fluorescence imaging

For immunofluorescence detection, hBMECs seeded on 1% collagen coated glass coverslips were infected with SPN (MOI ~ 25) as described above. At desired time points post infection, cells were washed with RPMI to remove extracellular SPN and fixed with ice chilled methanol for 10 min at −20°C. After blocking in 3% BSA for 2 h, cells were treated with appropriate primary antibody in 1% BSA for overnight at 4°C. Cells were then incubated with suitable secondary antibody in 1% BSA for 1 h at RT and finally, mounted with VectaShield, with or without DAPI (Vector Laboratories) for visualization using a Laser Scanning Confocal microscope (Zeiss LSM780) under 40X or 63X oil objectives. For live cell imaging, transfected hBMECs were grown on glass bottom petri dishes and infected with SPN as described earlier. The time lapse imaging was performed at multi-position under 63x oil immersion objective of Laser Scanning Confocal Microscope (LSM 780). The fixed and live images were acquired after optical sectioning and then processed using Zen lite software (Version 5.0).

### Western blot analysis

Following tetracycline treatment for 15 h, hBMECs were washed 2-3 times with chilled PBS and lysed in ice-cold RIPA buffer (50 mM Tris-Cl, pH 7.8, 150 mM NaCl, 1% Triton X-100, 0.5% Sodium deoxycholate, 1% SDS) containing protease inhibitor cocktail (Sigma Aldrich), Sodium fluoride (10 mM), PMSF (100 μM) and EDTA (5 mM). Total cellular proteins were separated from the cell debris by centrifugation (15000g for 30 mins at 4°C) of cell lysates. Proteins (10 μg or 20 μg) were separated on 12% SDS PAGE gels and were then transferred to activated PVDF membrane (Bio-Rad). Following 2 h of blocking in 5% skimmed milk, blot was incubated with appropriate primary and HRP-conjugated secondary antibodies and finally developed using ECL substrate (Bio-Rad).

### MTT cell viability assay

Effect of inhibitor and antibiotic treatments on the viability of hBMECs were quantified using EZcount™ MTT Cell Assay Kit as per manufacturer’s protocol (HiMedia). Briefly, treated hBMECs were washed with Hanks’s Balanced salt solution containing calcium and magnesium (HBSS-Ca^+2^ Mg^+2^). 500 μl of 10% MTT reagent in HBSS-Ca^+2^ Mg^+2^ from a stock of 5 mg/ml was added to cells and incubated at 37°C for 2 h. Further, cells were treated with 500 μl of solubilizing agent (10% Triton X-100, 90% isopropanol and 0.02% of 37% HCl) followed by repeated pipetting and incubation at RT for 30 min under shaking conditions to dissolve formazan crystals. Absorbance of the dissolved purple formazan crystals in the cell lysates was determined at 560 nm using a microplate spectrophotometer. Inhibitor non-treated infected and inhibitor non-treated infected cells were used as negative control while Triton X-100 (0.025%) was used as positive control.

### Statistics

All statistical analysis was performed using GraphPad Prism software (version 5) and significance level was set at *p* < 0.05. Statistical tests employed for individual experiments are mentioned in the respective figure legends. At least three independent experiments were performed. The results are presented as the mean ± SD of one representative experiment.

## ACKNOWLEDGEMENT

We acknowledge the Biosafety Level 2 facility and Confocal Microscopy facility at IIT Bombay. We also thank Dr. Ninad Mehendale from K.J. Somaiya College of Engineering, Mumbai, India, for assistance with the editing of time-lapse videos. SB acknowledges financial support from UGC, Govt. of India. AB acknowledges research funding from Science and Engineering Research Board (SERB), Govt. of India (Grant No. EMR/2016/005909). The funder had no role in study design, data collection and interpretation, or the decision to submit the work for publication.

---

**FIG S1. Cell viability, invasion and intracellular survival in presence of autophagy inhibitor 3-MA and proteasome inhibitor MG132.**

**A.** Cell viability of hBMECs following pre-treatment with autophagy inhibitor 3-MA (1 mM) or proteasome inhibitor MG132 (10 μM) was estimated by MTT assay. DMSO and Triton X-100 treated cells were used as negative and positive control, respectively. Results are expressed as percent cell viability with respect to negative control. Data is represented as mean ± SD of triplicate. Statistical analysis was performed using one-way ANOVA (Tukey’s multiple comparison test). ns, non-significant, *** p < 0.001.

**B.** Comparison of invasion efficiencies of SPN following treatment with 3-MA or MG132 or DMSO as vehicle. Bars represent mean ± SD. Statistical analysis was performed using one-way ANOVA (Dunnett’s multiple comparisons test). ns, non-significant.

**C. D.** Percent intracellular survival of internalized SPN in presence of 3-MA (1 mM) (C) or MG132 (10 μM) (D) at indicated time points relative to 0 h. Data are presented as mean ± SD of triplicate hBMEC cultures. Statistical analysis was performed using two-way ANOVA (Bonferroni test). ** p < 0.005, *** p< 0.001.

**FIG S2. Association of proteasome or autophagy with SPN depends on ubiquitin topology associated.**

**A, C.** Confocal micrographs showing no association of K48-Ub (red) positive SPN (blue) with LC3 (green) (A) or K63-Ub (magenta) marked SPN (blue) with LMP2 (green) (C) at 9 h p.i. hBMECs stably expressing LC3-GFP (A) or GFP-LMP2 (C) were infected with Hoechst stained SPN and stained with Anti-K63-Ub and Anti-K48-Ub linkage specific Abs. Arrowhead designates event shown in insets. Scale bar, 5 μm.

**B, D.** Fluorescent line scan across the yellow line in the merged inset in “A and C” is shown in “B and D”, respectively.

**FIG S3. Cell viability and invasion efficiency of ubiquitin variants expressing cells in presence of tetracycline.**

**A.** Viability of hBMECs expressing Ub^K48R^-GFP or mStrawberry-Ub^K63R^ or mStrawberry following treatment with tetracycline (40 μg/ml) as determined by MTT assay. DMSO and Triton X-100 treatments were used as negative and positive controls, respectively. Results are expressed as percent cell viability with respect to negative control. Bars are mean ± SD. Statistical analysis was performed using one-way ANOVA (Tukey’s multiple comparison test); ns, non-significant; *** p < 0.001.

**B.** Invasion efficiency of SPN in hBMECs expressing Ub^K48R^-GFP or mStrawberry-Ub^K63R^ or mStrawberry. Data are presented as mean ± SD of triplicate hBMEC cultures. Statistical analysis was performed using one-way ANOVA (Dunnett’s multiple comparisons test). ns, non-significant.

**S1 Movie. Proteasome dependent** c**learance of ubiquitinated SPN in brain endothelium.** Time lapse fluorescence imaging of DRAQ5 stained SPN’s (blue) degradation inside hBMECs stably expressing mStrawberry-Ub (red) and GFP-LMP2 (green). Live cell imaging was performed under laser scanning confocal microscope starting at 6 h p.i. The video depicts ubiquitinated SPN clearance while being associated with proteasome. Time interval, 30 min; scale bar, 1.5 μm.

**S2 Movie. Ubiquitinated SPN killing by autophagy.** Time lapse fluorescence imaging of DRAQ5 stained SPN’s (blue) degradation of inside hBMECs stably expressing mStrawberry-Ub (red) and LC3-GFP (green). Live cell imaging was performed under laser scanning confocal microscope starting at 6 h p.i. The video depicts killing of ubiquitinated SPN in an autophagy-dependent manner. Time interval, 10 min; scale bar, 1.5 μm.

