## Supplementary figures and images for "Differential ubiquitination as an effective strategy employed by Blood-Brain Barrier for prevention of bacterial transcytosis"

### Fig. S1

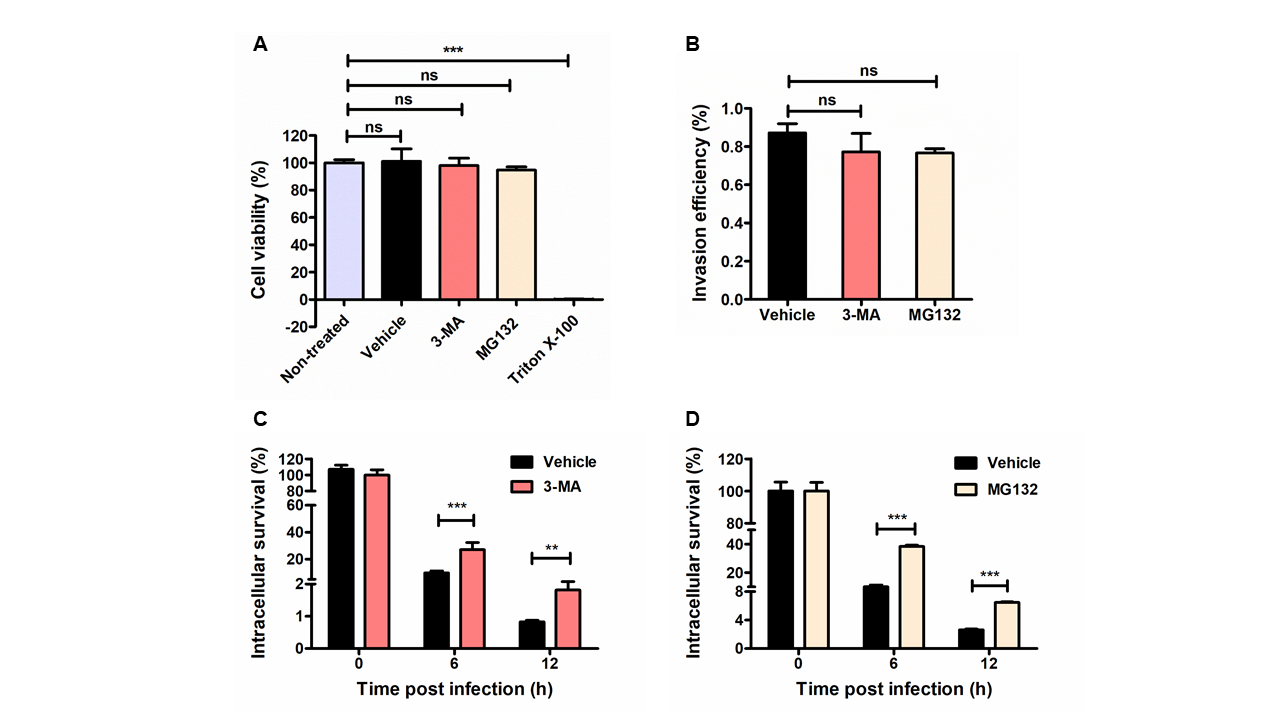

### Fig. S2

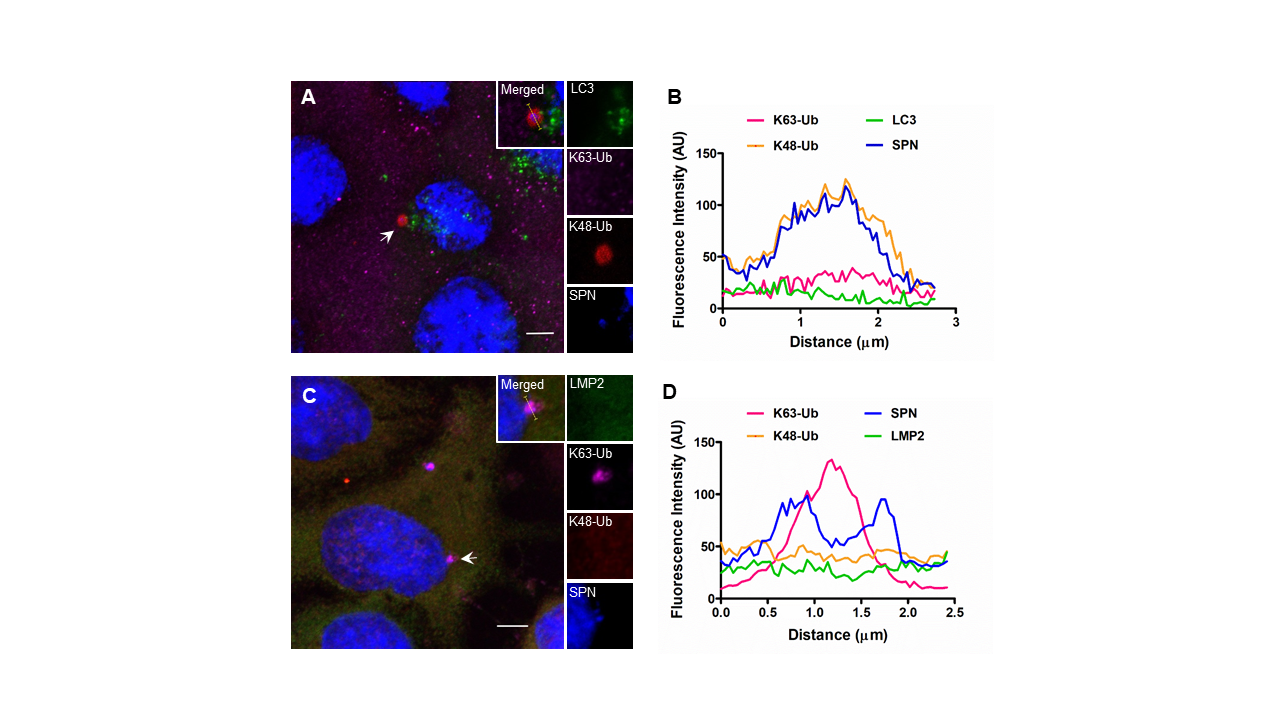

### Fig. S3

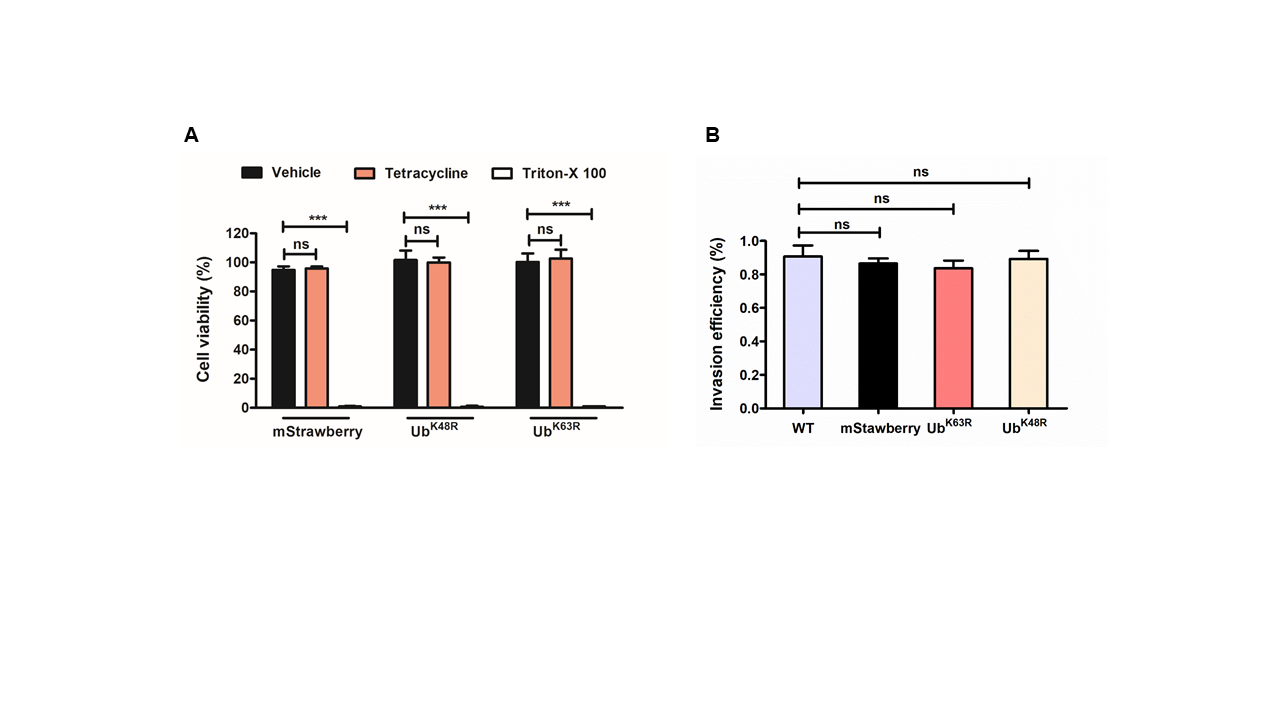
